# Life history evolution and phenotypic plasticity in parasitic eyebrights (*Euphrasia*, Orobanchaceae)

**DOI:** 10.1101/362400

**Authors:** Alex D. Twyford, Natacha Frachon, Edgar L. Y. Wong, Chris Metherell, Max R. Brown

## Abstract

**Premise of the study:** Parasite lifetime reproductive success is determined by both genetic variation and phenotypically plastic life history traits that respond to host quality and external environment. Here, we use the generalist parasitic plant genus *Euphrasia* to investigate life history trait variation, in particular whether there is a trade-off between growth and reproduction, and how life history traits are affected by host quality.

**Methods:** We perform a common garden experiment to evaluate life history trait differences between eleven *Euphrasia* taxa grown on a common host, document phenotypic plasticity when a single *Euphrasia* species is grown on eight different hosts, and relate our observations to trait differences recorded in the wild.

**Key results:** *Euphrasia* exhibit a range of life history strategies that differ between species that transition rapidly to flower at the expense of early season growth, and those that invest in vegetative growth and delay flowering. Many life history traits show extensive phenotypic plasticity in response to host quality and demonstrate the costs of attaching to a low-quality host.

**Conclusions:** Common garden experiments reveal trait differences between taxonomically complex *Euphrasia* species that are characterised by postglacial speciation and hybridisation. Our experiments suggest life history strategies in this generalist parasitic plant genus are the product of natural selection on traits related to growth and flowering. However, host quality may be a primary determinant of lifetime reproductive success.

## BACKGROUND

Parasitism is a ubiquitous feature of the natural world, with parasitic organisms present in every ecosystem and found to exploit all free-living organisms (Price, 1980; Windsor, 1998). The life history of parasitic organisms, in particular their developmental timing, must be synchronised with that of the host to maximise a parasite’s individual growth, survival and fecundity. However, the life history of external parasites (exoparasites) must not only be timed to their host, but to external environmental conditions (Fain, 1994). Changes in the developmental timing of a parasite will primarily be plastic when responding to local conditions or fluctuating environments, or genetic in response to more predictable or spatially structured variation (Reed et al., 2010). Understanding how genetic and plastic differences contribute to life history trait variation in parasite species and populations is essential for understanding parasite speciation and host-parasite interactions (Birget et al., 2017).

Parasitic plants are a group of c. 4500 species of at least 12 separate evolutionary origins that have evolved a modified feeding organ, the haustorium, which allows them to attach to a host plant and extract nutrients and other compounds (Westwood, Yoder, and Timko, 2010; Twyford, 2018). Parasitic plants are morphologically diverse and present a broad range of life history strategies (Schneeweiss, 2006; Těšitel, Plavcová, and Cameron, 2010) with the host species range a critical trait that determines parasite interactions and performance in the wild. Generalist parasitic plants can often attach to a broad range of hosts, with the well-studied grassland parasite *Rhinanthus* found to attach to over 50 co-occurring grass and herbaceous species (Cameron, Coats, and Seel, 2006). All generalist parasitic plants are exoparasites whose leaves, stems, roots and flowers grow outside the host and only the haustorium invades and grows within the host (Twyford, 2017).

The range of hosts and their visible external growth make hemiparasitic plants ideal for investigating life history trait evolution and host-parasite interactions. To date, research has largely focused on three aspects of life history variation in parasitic plants. Firstly, a body of research has looked to understand variation for specific traits between populations and related species. For example, work on the hemiparasite *Pedicularis* has shown how investment in male reproductive organs primarily depends on extrinsic environmental conditions rather than intrinsic resources (Guo, Mazer, and Du, 2010a), and how seed mass depends on intrinsic factors rather than elevation (Guo, Mazer, and Du, 2010b). Secondly, researchers have investigated how life history traits are affected by host quality. In the widespread and weedy obligate hemiparasite *Phelipanche ramosa*, the duration of the lifecycle differs between 14 weeks and 40 weeks depending on host (Gibot-Leclerc et al., 2013), with evidence of local host adaptation. Finally, a number of studies have looked at life history variation between species studied in a phylogenetic context (Schneeweiss, 2006; Těšitel et al., 2010). For example, broad-scale analyses of the Rhinantheae clade in the Orobanchaceae has shown a shift from a perennial ancestor to annuality, with correlated shifts to a reduced seed size (Těšitel et al., 2010). Despite the diversity of this research, there are still considerable gaps in our knowledge as to how life history trait variation is maintained (e.g. how common are trade-offs between life history traits), how much of this variation is genetic and how much is plastic, and which traits are the targets of natural selection.

In this study, we explore life history trait evolution in generalist parasitic eyebrights (*Euphrasia*, Orobanchaceae). *Euphrasia* is the second largest genus of parasitic plants with c. 350 species, and is characterised by recent transoceanic dispersal and rapid species radiations (Gussarova et al., 2008). In the United Kingdom, the 21 *Euphrasia* species demonstrate recent postglacial divergence with many species indistinguishable at DNA barcoding loci (Wang et al., 2018), with complex morphological variation (Yeo, 1968; Metherell and Rumsey, 2018), and found to readily hybridise (Stace, Preston, and Pearman, 2015). Despite this shallow species divergence, *Euphrasia* species demonstrates substantial ecological divergence, with many taxa restricted to specific habitats such as coastal turf, mountain scree or open grassland. Habitat differences would be expected to exert strong selection on life history traits, and this may include selection on growth to match seasonal water availability and to exploit local hosts, or selection on flowering time in response to local competition from surrounding plants, or in response to mowing or grazing.

Our research builds on a large body of experimental work with *Euphrasia*, which has been used in common garden studies for over 125 years (Koch, 1891). Early growth experiments revealed phenotypic differences observed in the field between the related species *E. rostkoviana* and *E. montana* are maintained in a common garden (Wettstein, 1895). Experimental work in the 1960s showed the growth of various *Euphrasia* species differs depending on the host species (Wilkins, 1963; Yeo, 1964). More recent replicated pot experiments of individual *Euphrasia* species or populations in a common garden (Matthies, 1998; Zopfi, 1998; Lammi, Siikamäki, and Salonen, 1999; Svensson and Carlsson, 2004) or in experimental field sites (Seel and Press, 1993; Hellström et al., 2004), have shown the effect of commonly encountered hosts such as grasses and legumes on parasite biomass, mineral accumulation, plant architecture and reproductive output. Despite this extensive experimental work, studies in *Euphrasia* have yet to compare life history strategies of different species, and the extent of phenotypic plasticity in life history traits. This work is critical for understanding the nature of species differences in a taxonomically complex group, as well as for evolutionary studies of parasite diversity. It also unclear whether *Euphrasia* are restricted to growing on hosts such as grasses and herbaceous species, or can parasitize a broad range of taxa including novel hosts rarely encountered in the wild. To address these questions requires simultaneously investigating the growth of multiple *Euphrasia* species and multiple host species with replication to enable suitable statistical comparisons.

Our experiments first assess the extent of life history trait variation among several *Euphrasia* species and their hybrids when grown on a single host species in standardised common garden conditions. This experiment addresses whether there is life history trait divergence among recently diverged parasite species. We then inspect plasticity of a single focal *Euphrasia* population grown on many different hosts. This experiment tests whether they are truly generalist parasites by growing them on a wide range of hosts as well as growing them without a host. Finally, we relate our trait observations made in a common garden to recordings made on herbarium specimens collected in the wild. This comparison will help us understand whether life history trait differences hold in both the common garden and in nature. Overall, our joint observations of phenotypic variation between closely related taxa, and the extent of host-induced plasticity within a species, both in an experiment and in the wild, provide new insights into life history variation in parasitic taxa. These results provide valuable comparisons to other generalist parasitic plants such as *Rhinanthus* and *Melampyrum*, to see similarities in life history across grassland parasitic plants.

## MATERIALS AND METHODS

### Experimental design and plant cultivation

We performed two common garden experiments to investigate phenotypic variation in parasitic *Euphrasia*. Our species differences experiment observed life history trait differences of twenty four populations from five species and six natural hybrids when grown on clover (*Trifolium repens*), which is often used as a host in parasitic plant experiments because the parasite grows vigorously and has high survival (Zopfi, 1998). This experiment includes multiple populations of three widespread and closely related species, *E. arctica, E. confusa* and *E. nemorosa*, and sparse population sampling of two more distinct taxa, *E. micrantha* (one population) and *E. pseudokerneri* (two populations). Our phenotypic plasticity experiment assessed the impact of eight potential hosts (*Arabidopsis thaliana, Equisetum arvense, Festuca rubra, Holcus lanatus, Marchantia polymorpha, Pinus sylvestris, Plantago lanceolata*, and *Trifolium repens*), and growing without a host, on the phenotype of a single focal taxon, *E. arctica*. These hosts were chosen to represent a broad representation of functional groups and phylogenetic diversity, with species encountered in the wild as well as novel hosts (full details in Table S1). The novel hosts were included to see the limits to which parasitic *Euphrasia* can benefit, namely with a woody tree (*Pinus*), a pteridophyte that produces adventitious roots (*Equisetum*), and a liverwort that produces rhizoids (*Marchantia*). Both common garden experiments took place in parallel in 2016. Wild-collected open-pollinated *Euphrasia* seeds were contributed by plant recorders as part of the ‘Eye for Eyebrights’ (E4E) public engagement project and as such included a scattered geographic sample across Great Britain (Table S2). Species were identified from herbarium specimens by *Euphrasia* referee Chris Metherell. Host seeds were sourced from commercial suppliers and from wild-collected material (Table S1).

Reliable cultivation of *Euphrasia* can be challenging due to low seed germination, variation in time to establishment, the requirement of seed stratification, and high seedling mortality when transplanted (Yeo, 1961; Zopfi, 1998). Here, we develop cultivation protocols that combine winter germination cues that improve germination and mimic nature, but also use highly standardised and replicated pot conditions that avoid transplanting *Euphrasia* and thus maximise survivorship. We planted a single *Euphrasia* seed into 9 cm plastic pots filled with Melcourt Sylvamix Special growing media in December, with plants left outside overwinter to experience natural seed stratification. Hosts were planted in seed trays in April. *Euphrasia* plants were moved to an unheated and well-ventilated greenhouse at the Royal Botanic Garden Edinburgh (RBGE) in the spring once the cotyledons were fully expanded, and a single seedling from each host (or a 1cm^2^ clump of *Marchantia*) introduced. Hosts that died within ten days of planting were replaced. Twenty or more replicates were grown for each host-parasite combination. Plants were subsequently grown to flowering with regular watering, randomised at weekly intervals, and foreign weed seedlings removed.

### Trait measurements and statistical analyses

We measured a suite of seven morphological traits at first flowering. These traits are characters related to life history variation, indicators of plant vigour, or characters used in taxonomy. In addition to date of first flowering, we measured: corolla length, cauline leaf length:internode length ratio (‘internode ratio’), number of leaf teeth on the lower floral leaf, number of nodes to flower, number of branches and plant height. All length measurements were made to the nearest millimetre, and followed Metherell and Rumsey (2018). For the phenotypic plasticity experiment, we also recorded early season growth (height six weeks after transplantation of potential host) and height at the end of season after senescence. We did not make direct observations of below-ground attachment due to the fine root structure of *Euphrasia*. Instead, we inferred that attachment is likely to have taken place based on observations of height, following Yeo (1964).

We analysed data for the two experiments separately but with similar statistical approaches. For the species differences experiment we used generalised linear mixed effects models to fit species as a fixed effect and population as a random effect. Population was implemented as a random effect because we were interested in the variability between populations, as well as to account for it for estimating species parameters. For the phenotypic plasticity experiment we used generalised linear mixed effects models with presence or absence of host as a fixed effect and host species identity as a random effect. We used a Gaussian distribution for all models except where the response were positive integer counts, where we used a Poisson distribution. To calculate the significance of a fixed or random effect we used Likelihood Ratio Tests where the full model was compared with a nested model where one effect had been dropped. When a random effect term was dropped for each variable, we implemented generalised linear models using R base functions. For the trait number of branches the generalised linear mixed effects models failed to converge so we used MCMCglmm (Hadfield, 2010), with the best fitting of either a full or nested model chosen based on a lower Deviance Information Criterion (DIC) value. For the phenotypic plasticity experiment, models for number of branches failed to converge in every model run, so this model was omitted. Using the flexible variance structures accommodated by MCMCglmm we were able to calculate “variance explained” by random effects in each model. In the case of a Gaussian model, variance explained by an explanatory variable is the fraction of the total variance accounted for by the variable. In the case of a Poisson model the calculation is more complex due to the mean-variance relationship (Nakagawa and Schielzeth, 2010). The models were run for 91,000 iterations with a burn-in of 21,000 and a thinning interval of 70. The priors were standard inverse-Wishart for the (co)variances and parameter expanded where there was poor mixing. Overdispersion was accounted for in each case by either implementing quasipoisson models or in mixed effect models by implementing an observation level random effect. We calculated r^2^ values from Pearson Correlation Coefficients between all traits and used Principal Component Analyses to quantify phenotypic clustering of individuals. All analyses were done in R version 3.4.3, with the packages lme4 (Bates et al., 2014) and MCMCglmm for generalised linear mixed effects models, Hmisc for correlations (Harrell Jr and Harrell Jr, 2018) and ggplot2 for data visualisation (Roff, 1992).

### Trait variation in the wild

We tested how phenotypes in the experiments relate to phenotypes in nature, by comparing results from the species differences experiment to phenotypic measurements made on herbarium specimens of the same populations sampled in the wild. Pressed plants submitted by field collectors varied in quality, and these sometimes missed the base of the plant, therefore we were unable to measure height. We measured traits on three individuals from each collection sheet for a given population. We used generalised linear mixed effects models to fit traits as a response variable, with where individuals were grown (i.e. common garden or wild-collected) as an explanatory variable. This two-levelled categorical factor was implemented as a fixed effect. Adding species as a random effect shows how much of the variability of the trait was explained by species given where individuals were measured. Count data were analysed with a Poisson distribution, in all other cases a Gaussian distribution was used. R-values were calculated using Pearson’s correlations of the population level means between the common garden and the wild samples. Analyses were performed in MCMCglmm.

## RESULTS

### Species differences

Our species differences experiment reveals extensive morphological trait variation across *Euphrasia* species when compared at first flowering. From the 222 individuals surviving to flowering on their clover host, we see a 3-fold difference in mean height, while other traits also proved variable (Fig. 1eh; Table S3). A large degree of this variation is separated by species and by population (Table 1). The species with the most distinct lift-history strategy is *E. micrantha*, which rapidly transitions to flower from a low node on the plant (8.3 ± 0.2 nodes), and flowers while it is short (70.6 mm ± 8.1). It also forms a distinct cluster in the PCA analysis (Fig. S1). *E. pseudokerneri* is relatively distinct, flowering once it has grown tall (176.4 mm ± 15.6) and from a high node on the plant (13.2 ± 0.4 nodes), but shows little separation in the PCA analysis. The morphologically similar *E. arctica, E. confusa* and E. *nemorosa* are partly distinct, with *E. nemorosa* initiating flowering 14 days later and from 3.3 nodes higher than *E. arctica*, but overlaps in many other traits and in overall multi-trait phenotype (Fig. S1). In most cases hybrids combine morphological characters of their parental progenitors, for example hybrids involving *E. nemorosa* flower later in the season and initiate flowering from a higher node than *E. arctica* hybrids (Fig. 1a-d).

**Figure 1.**
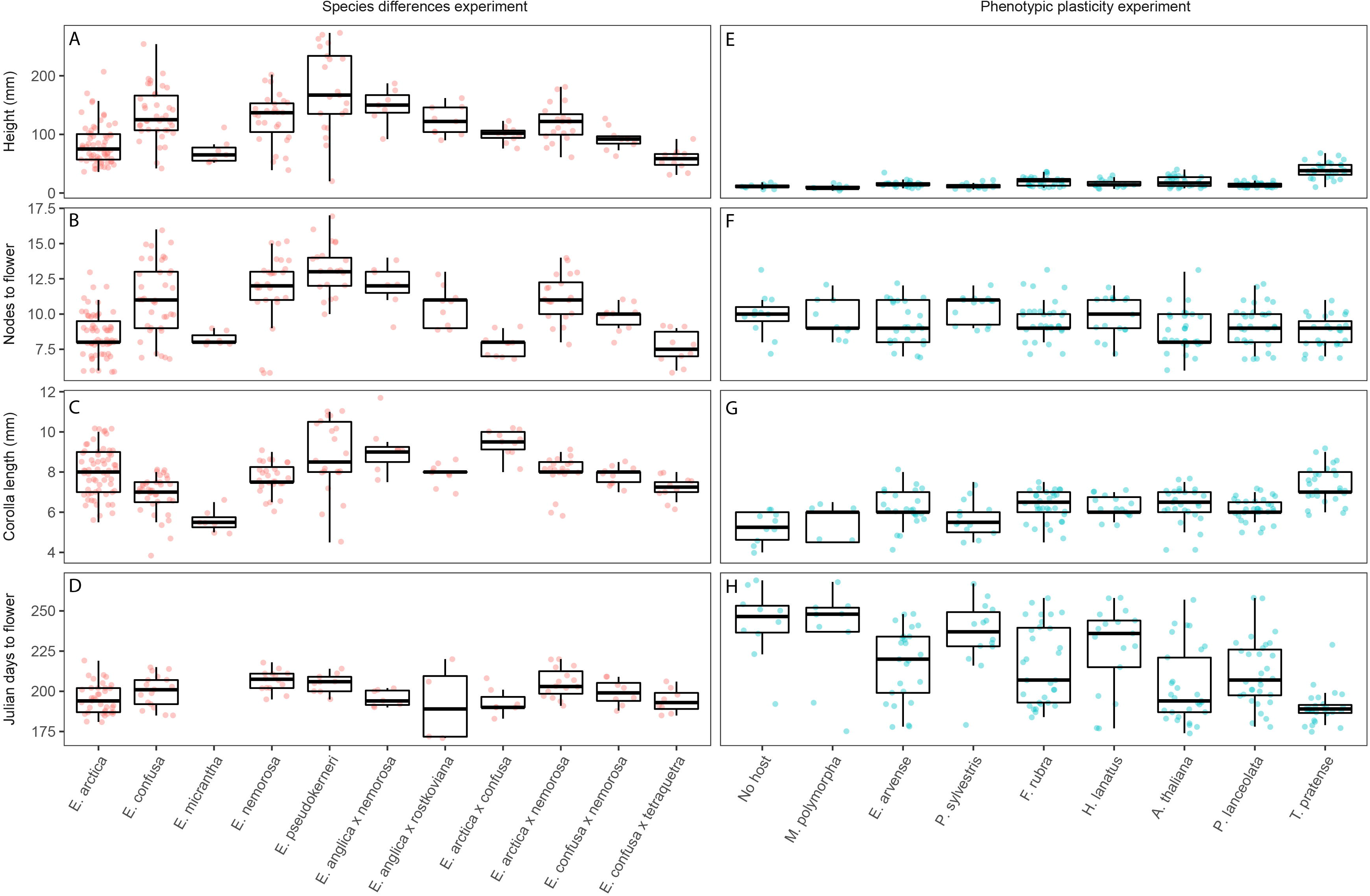
Trait variation in a common garden experiment of diverse *Euphrasia* species and hybrids grown on clover (A-D) and *E. arctica* grown on many different hosts (E-H). The edges of the boxplots show the first and third quartiles, the solid lines the median, the whiskers the highest and lowest values within 1.5-fold of the inter-quartile range and the dots each individual measurement.

**Table 1.** Summary of generalised linear mixed effects models of the influences on life history, plant vigour, and taxonomic traits for *Euphrasia* in a common garden. Table summarises model outputs for the species differences and the phenotypic plasticity experiment. The percentage variance explained by random effects are reported in brackets along with the 95% credibility interval. fModels for number of branches were implemented with a different statistical approach in MCMCglmm (see methods). *** P < 0.001, ** P < 0.01, * P < 0.05

|  | Species differences |  | Phenotypic plasticity |  |
| --- | --- | --- | --- | --- |
|  | Species | Population | Host/no host | Host |
|  |  | DIC<sub>full</sub> = 867.36; |  |  |
| Branches<sup>†</sup> | DIC<sub>full</sub> = 867.36; | DIC<sub>pop</sub> = 905.68 |  |  |
|  | DIC<sub>spp</sub> = 869.11 | (27.7%, 16.4 - 61.2%) | n/a | n/a |
| | | $\chi^2(1) = 42.89^{***}$ | | $\chi^2(1) = 28.79^{***}$ |
| Corolla length | $\chi^2(1) = 22.38^*$ | (49.1%, 28.8 - 75.5%) | $\chi^2(1) = 3.19$ | (13.1%, 5.6 - 52.3%) |
| | | $\chi^2(1) = 67.68^{***}$ | | $\chi^2(1) = 91.15^{***}$ |
| Height | $\chi^2(1) = 20.10^*$ | (57.2%, 37.3 - 80.2%) | $\chi^2(1) = 0.9033$ | (54.1%, 35.9 - 85.5%) |
| | | | | $\chi^2(1) = 6.94^{**}$ |
| Internode ratio | $\chi^2(1) = 24.5^{**}$ | $\chi^2(1) = 27.39^{***}$ | $\chi^2(1) = 1.38$ | (4.6%, 0.8 - 23.0%) |
| Julian days to | | $\chi^2(1) = 1.34$ | | $\chi^2(1) = 205.2^{***}$ |
| flower | $\chi^2(1) = 8.34$ | (21.2%, 7.7 - 64.6%) | $\chi^2(1) = 2.34$ | (37.4%, 13.4 - 66.3%) |
| | | $\chi^2(1) = 0.71$ | | $\chi^2(1) = 0$ |
| Node to flower | $\chi^2(1) = 26.66^{**}$ | (10.1%, 0.4 - 31.6%) | $\chi^2(1) = 0.38$ | (1%, 0.18 - 9.8%) |
| Number of leaf |  |  |  |  |
| teeth | $\chi^2(1) = 23.43^*$ | $\chi^2(1) = 0$ | $\chi^2(1) = 1.49$ | $\chi^2(1) = 5.46^*$ |

Correlation analyses across species reveal clear suites of traits that are related. Significant correlations were found between 14 of the 21 pairwise comparisons, with 5 of these correlations with an R > 0.6 (Table 2a). Plants flowering at a late node are more likely to be tall, more highly branched, as well as having many teeth on the lower floral leaf. The relationship of traits is also supported in the PCA analysis, with many traits contributing to multiple principal components (Table S4). These height- and flowering node-related traits are largely uncorrelated with cauline internode-leaf ratio and corolla length.

**Table 2.**
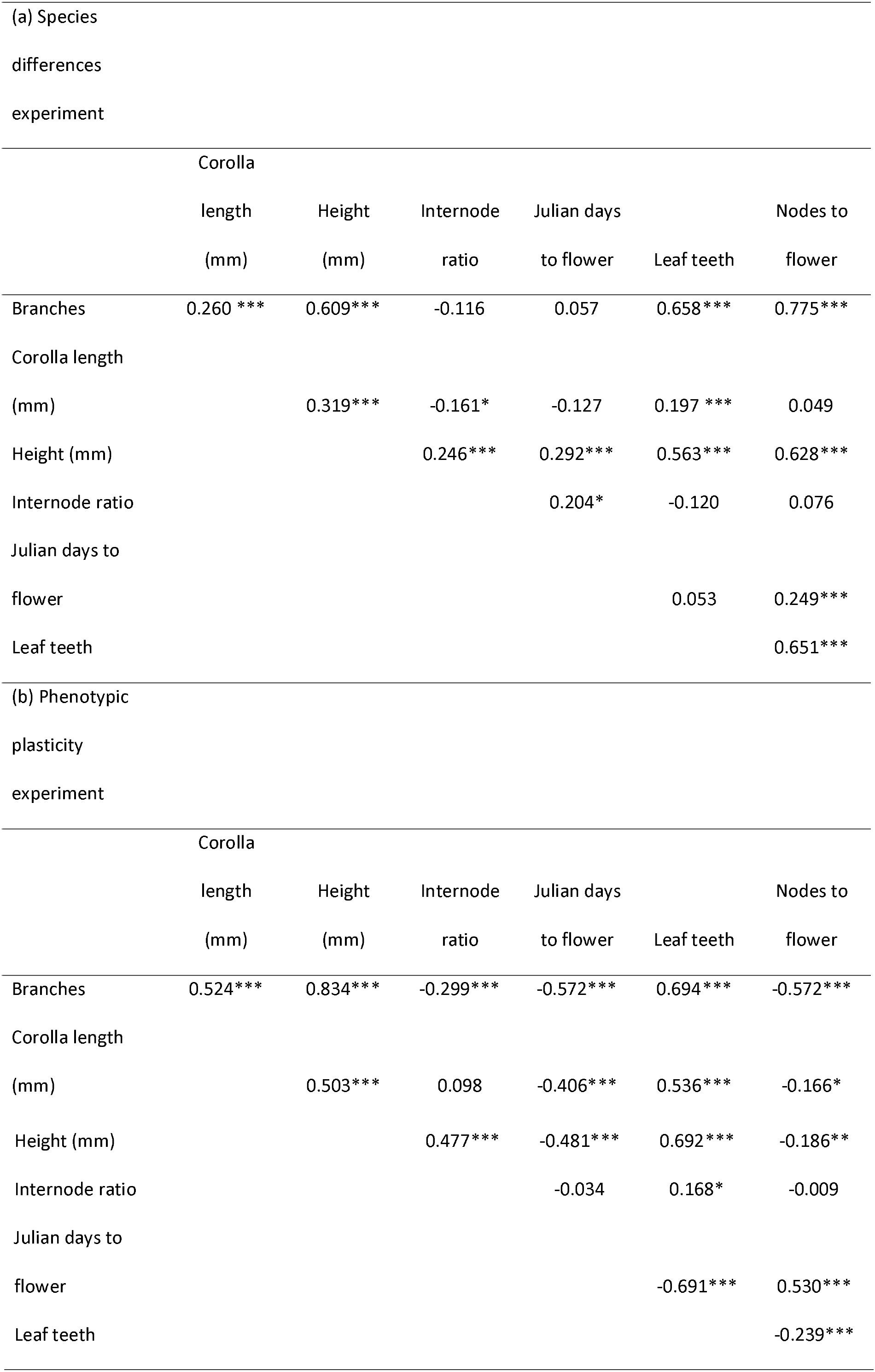
Pearson’s correlation coefficients for seven phenotypic traits measured in a common garden experiment for (a) Five *Euphrasia* species and 6 hybrids, (b) *Euphrasia arctica* grown with 8 hosts and without a host. *** P < 0.001, ** P < 0.01, * P < 0.05. Asymptotic p-values values are reported from the Hmisc package in R using the rcorr() function.

### Phenotypic plasticity

Our phenotypic plasticity experiment shows substantial morphological variation across 194 *E. arctica* plants grown with 8 different host species, and the 22 plants grown without a host. Plants growing on clover transitioned to flower quickly, grew tall by the time of first flowering, and produced large flowers (Fig. 1e-h, Table S5). This contrasts with *Euphrasia* with no host, which flowered on average 53 days later, were extremely short at first flowering, and produced small flowers. Despite this, the presence or absence of host was not significant (Table 1), due to *Euphrasia* grown with *Marchantia* and *Pinus* growing similarly to the no host treatment (Fig. 1e-h). The other hosts, *Arabidopsis, Equisetum, Festuca, Holcus* and *Plantago* conferred some benefit for *Euphrasia* and fell between clover and no host. Our comparison of growth across host treatments measured through the year showed that height at the end of the season is weakly predicted from height 6-weeks after introducing a host (R = 0.47), but strongly correlated with height at first flowering (R = 0.82; Fig. S2). Plants that flowered early were more likely to grow larger by the end of season (R = -0.55) and become more highly branched (R = -0.57).

Across host treatments, there was a significant negative correlation between Julian days to flower and most other traits (Table 2b). We find that late flowering individuals are likely to be smaller at first flowering, are less highly branched, have less highly toothed leaves, and have smaller flowers (Table S5). While these traits were strongly correlated, there were substantial differences in the amount each trait changed. Days to flower differed considerably depending on host, with a 3.8x greater difference than seen between means of different *Euphrasia* species (Fig. 1d & 1h). In contrast corolla length and node to flower proved less variable depending on host, with a 1.4x and 1.2x change between means, respectively.

### Variation in the wild

The comparison between the species differences common garden experiment and wild-collected herbarium specimens revealed a single trait, nodes to flower, is strongly correlated (R=0.79) and not significantly different (p_MCMC_=0.71) between environments (Fig. 2). All other traits did differ significantly between environments (p_MCMC_ < 0.05), with *Euphrasia* plants in the common garden having corollas on average 1.4mm longer, with 0.2 more teeth on the lower floral leaves, an increase in cauline:internode ratio of 1.0mm, and 4 more pairs of branches. Despite differing between environments, trait values were highly correlated for corolla length (R= 0.93, p_MCMC_ < 0.001) and cauline internode:leaf length ratio (R=0.65, p_MCMC_ < 0.001), but not for number of leaf teeth (R=0.07, p_MCMC_ = 0.034) and number of branches (R=0.29, p_MCMC_ < 0.001).

**Figure 2.**
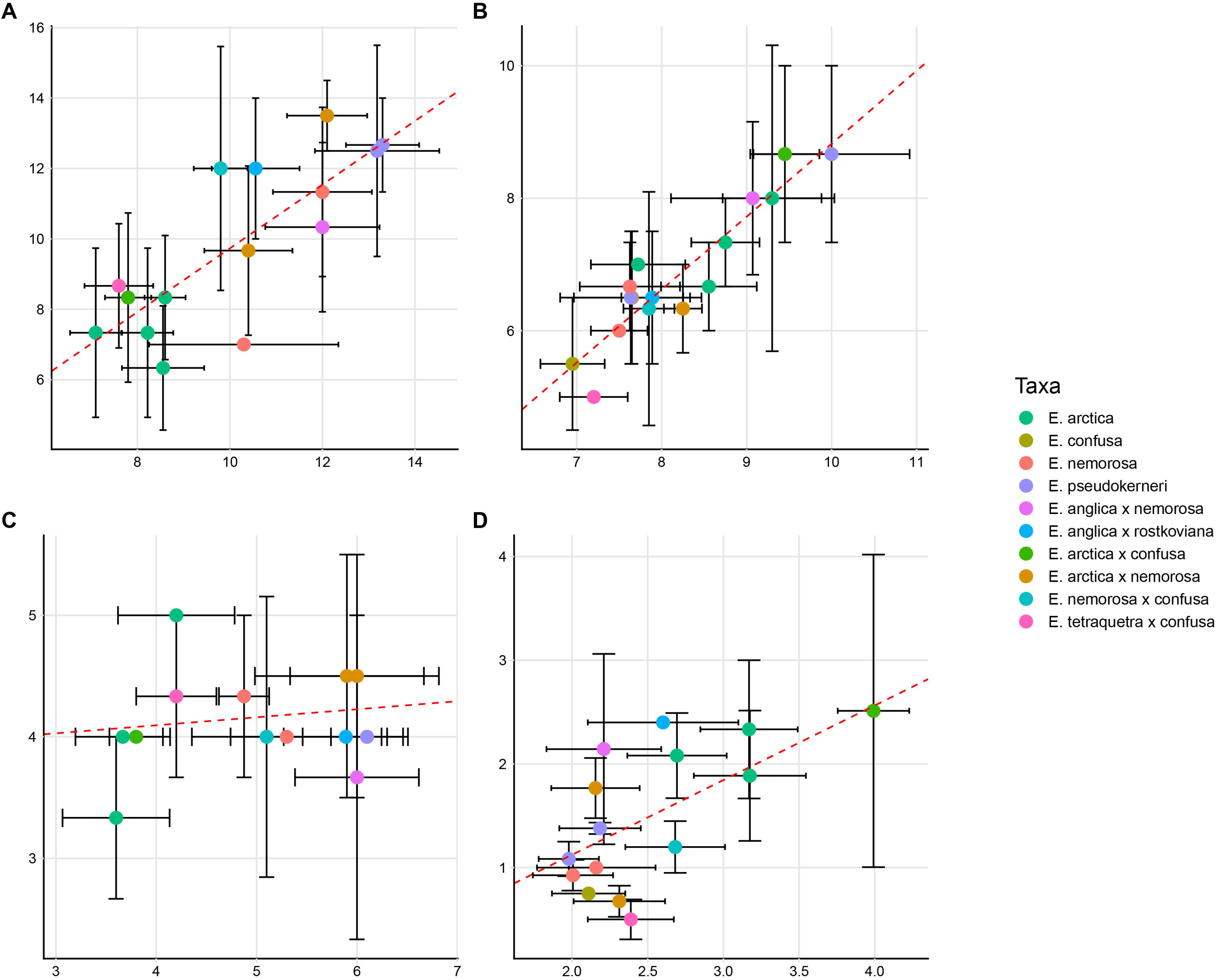
Relationship between morphological trait measurements made in the common garden and on wild-collected herbarium specimens for (A) nodes to flower, (B) corolla length (mm), (C) number of leaf teeth, (D) cauline leaf:internode ratio. Points are for populations, with bars representing the standard error of measurements. The line of best fit was calculated using coefficients from linear regression models on the means of each *Euphrasia* population.

## DISCUSSION

Our study sheds new light on the life history evolution and phenotypic plasticity of the generalist parasitic plant *Euphrasia*. We find different life history strategies between recently divergent species, with some species rapidly transitioning to flower at the expense of growth-related traits, while others delay flowering and invest in early-season vegetative growth. However, many traits related to life history differences are phenotypically plastic and show major changes in response to host quality. In addition to our experimental observations, the consistency between life history traits in a common garden and in the wild suggest our experimental observations generalise to patterns observed in nature, and that currently delimited *Euphrasia* species are, at least in part, distinct. Overall our study highlights the value in integrating multiple common garden experiments and field collections to study life history strategies in parasitic plants, and demonstrates the rapid evolution of life history differences in a postglacial radiation of parasites. We contrast our results with other well-studied hemiparasitic taxa to look for general patterns of life history evolution in grassland parasites.

### Life history variation in a parasitic plant

Our study finds evidence for different life history strategies between *Euphrasia* species in Britain. *E. arctica, E. micrantha* and hybrids such as *E. arctica* x *E. confusa*, transition rapidly to flower, flower while they are short, and produce their first flower from a low node on the plant. This contrasts with *E. pseudokerneri, E. nemorosa* and hybrids involving *E. nemorosa* that delay flowering until later in the season, grow tall before flowering, and produce their first flower from a late node on the main axis. These different life history strategies correspond to the known of ecology of these species, with *E. nemorosa* flowering late in tall mixed grassland, while *E. micrantha* flowers early in patchy heathland where competition is less intense (Metherell and Rumsey, 2018). A similar relationship between internode number and habitat have been observed in populations of *E. rostkoviana* in Sweden, where plants from intensely grazed pasture flower at a lower node in a common garden (Zopfi, 1998). Overall, these observations within and between populations are consistent with the classic life history trade-off between growth and reproduction (Roff, 1992; Stearns, 1992). For *Euphrasia* growing in the wild, early reproduction allows the plants to reliably complete their lifecycle before summer competition, herbivory, mowing, summer drought and other seasonal abiotic and biotic stresses. However early flowering involves reproducing at the expense of early season growth and at a time when the resource budget is constrained by relatively few haustorial connections. These trait trade-offs pose an interesting comparison to the well-studied *Mimulus guttatus (syn. Erythranthe guttata*), a non-parasitic relative in the Lamiales that shares the same basic plant architecture. In *M. guttatus* multiple traits related to growth and reproduction are correlated both within and between populations, due to genetic trade-offs between time to flower and fecundity (Mojica et al., 2012; Friedman et al., 2015). In *Euphrasia* the genetics underpinning this life history trade-off have yet to be characterised, and may be a consequence of multiple independent loci or trade-offs at individual loci (Hall, Lowry, and Willis, 2010).

While much life history variation is captured by differences in time to flower and growth-related traits, we also see evidence for flower size representing a separate axis of variation across *Euphrasia* species. In our common garden *E. micrantha* has small corollas, while *E. arctica* and *E. nemorosa* have larger corollas, and corolla size is not strongly correlated with other traits. *Euphrasia* species are well-known to have flower size variation, with a continuum between small flowered species that are highly selfing (e.g. *E. micrantha*, corolla size = 4.5 – 6.5mm, inbreeding coefficient F_IS_ > 0.88; Stone, 2013), and large flowered species that are highly outcrossing (e.g. *E. rostkoviana* flower size 8 – 12mm, F_IS_ = 0.17 – 0.25; French et al., 2005). Such continuous variation in outcrossing rate has been documented in species of *Datura* (Motten and Stone, 2000), *Mimulus* (Karron et al., 1997) and *Nicotiana* (Breese, 1959). Small flowers have shorter anther-stigma separation and thus increased potential for autogamous selfing (Karron et al., 1997), while also having reduced attractiveness to pollinators and thus receiving less outcross pollen (Mitchell et al., 2004). In addition to differences in corolla size between *Euphrasia* species, corolla size also shows a change of up to two millimetres in response to host quality, with this change of a magnitude that may potentially affect the mating system (Luo and Widmer, 2013). This suggests host quality represents a previously unaccounted factor affecting the mating system of parasitic plants.

Our comparisons of *Euphrasia* species in a common garden also sheds light on the distinctiveness of these recently divergent species. *Euphrasia* is one of the most taxonomically complex plant genera, with the 21 currently described British species presenting complex and often overlapping morphological variation (French et al., 2008; Metherell and Rumsey, 2018; Wang et al., 2018). Our study suggests varying degrees of morphological distinctiveness of *Euphrasia* species. We see *E. micrantha* is clearly morphologically distinct and *E. pseudokerneri* mostly distinct, while the closely related species *E. arctica, E. confusa* and *E. nemorosa* differ in life history traits such as nodes to flower, but overlap in many other traits and are not clearly separated in the PCA. Moreover, our study is likely to overestimate the distinctiveness of taxa by only including a subset of UK species and by choosing populations that could be identified to species-level in the field. We suspect adaptive divergence between closely related *E. arctica, E. confusa* and *E. nemorosa* is a consequence of differential natural selection for local ecological conditions such as soil water availability or mowing. Selection appears to be operating at a fine spatial scale, with significant life history trait differences evident between populations within species. These taxa may be genetically cohesive, either showing genome-wide divergence, or divergence in genomic regions underlying life history differences (Twyford and Friedman, 2015). Alternatively, these taxa may be polytopic and not genetically cohesive (Hollingsworth, Neaves, and Twyford, 2017). Genomic sequencing of natural populations is hoped to resolve this issue.

### Phenotypic plasticity in response to host quality

Our phenotypic plasticity experiment shows *Euphrasia* benefit from growing with a range of hosts. Specifically, *E. arctica* with a high quality host such as clover rapidly transitions to flowering. Differences in growth are established early in the season, and early flowering plants go on to grow the tallest, are more highly branched and have the potential to produce many extra flowers. At the other extreme, growing with a poor host or without a host results in late-flowering minimally vegetative plants. Most hosts fall between these extremes and offer moderate return to the parasite. Two surprising results were that *E. arctica* parasitizing *Arabidopsis* grew relatively tall despite the host senescing early in the growth season, and that *Euphrasia* on *Equisetum* performed similarly to the commonly encountered grass *Holcus lanatus*, suggesting surprising attachment or indirect benefits through association with *Equisetum* fungal symbionts (Bouwmeester et al., 2007). Differences in host quality are complex, but may be attributed to root architecture, germination time and resource availability, as well as the presence of mechanisms to defend against parasite attack, such as cell wall thickening, localised host dieback, and chemical defence (Cameron, Coats, and Seel, 2006; Twyford, 2018). Many of these attributes of host quality appear to be shared between related hemiparasites in the Orobanchaceae, though contrasting results from *Rhinanthus* suggest there are not universal ‘good’ hosts for all parasitic plants (Matthies, 2017). While *Euphrasia* is generally thought to have low reliance on host resources, deriving only ~30% carbon heterotrophically (Těšitel, Plavcová, and Cameron, 2010), at least under our experimental conditions a good host is required in order to produce multiple flowers. Overall our results point to *E. arctica* being a true generalist parasite, but one that only grows better with a subset of hosts that it may encounter.

We observe considerable variation in the plasticity of traits in response to host quality. Only three pairs of trait correlations are consistent across both *Euphrasia* common garden experiments (between height, number of branches and leaf teeth), with the positive correlation between node to flower, height, and number of branches seen between species breaking down when *Euphrasia* are grown on different hosts. The most notable plasticity is seen in flowering time, with plants on the best hosts rapidly transitioning to flower within ~100 days of germination, while plants with a more typical host (e.g. *Holcus lanatus*) flower a month later. Phenotypic plasticity in flowering time in response to resource availability is well documented in many plant groups, particularly *Arabidopsis* (e.g. Zhang and Lechowicz, 1994), but has received less attention in studies of parasitic plants, which are more likely to look at growth-related traits such as biomass (Ahonen, Puustinen, and Mutikainen, 2005; Matthies, 2017). However date of first flowering has been show to differ by up to 10 weeks in populations of *Rhinanthus glacialis* across Switzerland (Zopfi, 1995). Overall we expect date of first flowering to be critical for the life time reproductive success of parasitic plants in the wild.

In contrast to traits showing extensive plasticity, we also see evidence of developmental constraint in number of nodes to flower. This trait showed the least plasticity with different hosts, is consistent between populations within species, and between the common garden and the field. This suggests that the developmental event of transitioning to flower is genetically determined, with changes in flowering time altered by plasticity in internode length and not nodes to flower. This may explain why nodes to flower is such an important diagnostic trait for species identification in *Euphrasia* and related species in the Rhinantheae (Jonstrup, Hedrén, and Andersson, 2016). Despite nodes to flower changing little in response to host quality, our overall impression is that *Euphrasia* show considerable plasticity and little developmental constraint in many aspects of growth, or that trait values are affected by complex interactions related to host-parasite attachment. In particular, the variation between individuals on a given host, suggests other variation, such as genetic background in host and parasite, as well as the timing of attachment, may be crucial in determining performance.

## CONCLUSIONS

Despite over a century of experimental studies in parasitic plants, our understanding of the evolution of life history strategies in these diverse organisms is extremely limited. Our results in *Euphrasia* provide strong support for the rapid evolution of distinct life history strategies in response to local ecological conditions, with phenotypic plasticity further altering plant growth in response to host availability. We anticipate that future studies that test life time reproductive success of many parasitic plant species grown on many different host species will give further insight into the complex nature of host-parasite interactions.

## Data accessibility

Phenotypic data from both common garden experiments and from herbarium collections, as well as the R scripts used for data analysis, are deposited in Dryad XXXXXX (accession to be given on acceptance).

## Acknowledgements

We are indebted to the *Euphrasia* research community and to botanical recorders for providing seed and herbarium specimens from natural populations. We also thank horticulture staff at the RBGE for help with plant care. Jarrod Hadfield provided statistical advice and Susanne Wicke provided useful information about parasitic plant biology. A.D.T. is supported by a Natural Environment Research Council (NERC) Fellowship NE/L011336/1 and grant NE/R010609/1. N.F. acknowledges funding from the Scottish Government’s Rural and Environment Science and Analytical Services Division (RESAS). M. B. is funded by BBSRC EASTBIO studentship.

## Author contributions

A.D.T conceived and designed the research. A.D.T., N.F. and E. L. Y.W. carried out the experiments. C.M. identified the plants. A.D.T. and M.B. analysed the data. A.D.T. and M.B. wrote the manuscript. All authors read and approved the manuscript.

## Data accessibility

R scripts for analysis are included in the Online Supporting Information. All phenotypic data are available from Dryad <Details given on acceptance>.

## Supporting information

**Figure S1.** Principal components analysis of morphological variation of *Euphrasia* in a common garden for (A) five species and six hybrids, (B) five species, (C) *Euphrasia arctica* with nine host treatments.

**Figure S2.** Relationship between early and midseason growth and flowering related traits and end of season height. (A) Height at first flowering, (B) height 6-weeks after germination, (C) Julian days to flower, (D) number of branches. Length measurements are reported in mm.

**Table S1.** Species and collection details for hosts used in the common garden experiment.

**Table S2.** Collection details for *Euphrasia* species used in the common garden experiment.

**Table S3.** Summary of trait values for many *Euphrasia* species and hybrids grown on a clover host. Values are mean +/- one standard error. Length measurements are in mm.

**Table S4.** Factor loadings for the principal components analyses of (A) five species and six hybrids, (B) five species, (C) *Euphrasia arctica* with nine host treatments.

**Table S5.** Summary of trait values for *Euphrasia arctica* grown on many different hosts. Values are mean +/- one standard error. Length measurements are in mm.

