## Supplementary figures and images for "Life history evolution and phenotypic plasticity in parasitic eyebrights (*Euphrasia*, Orobanchaceae)"

### Fig. S1

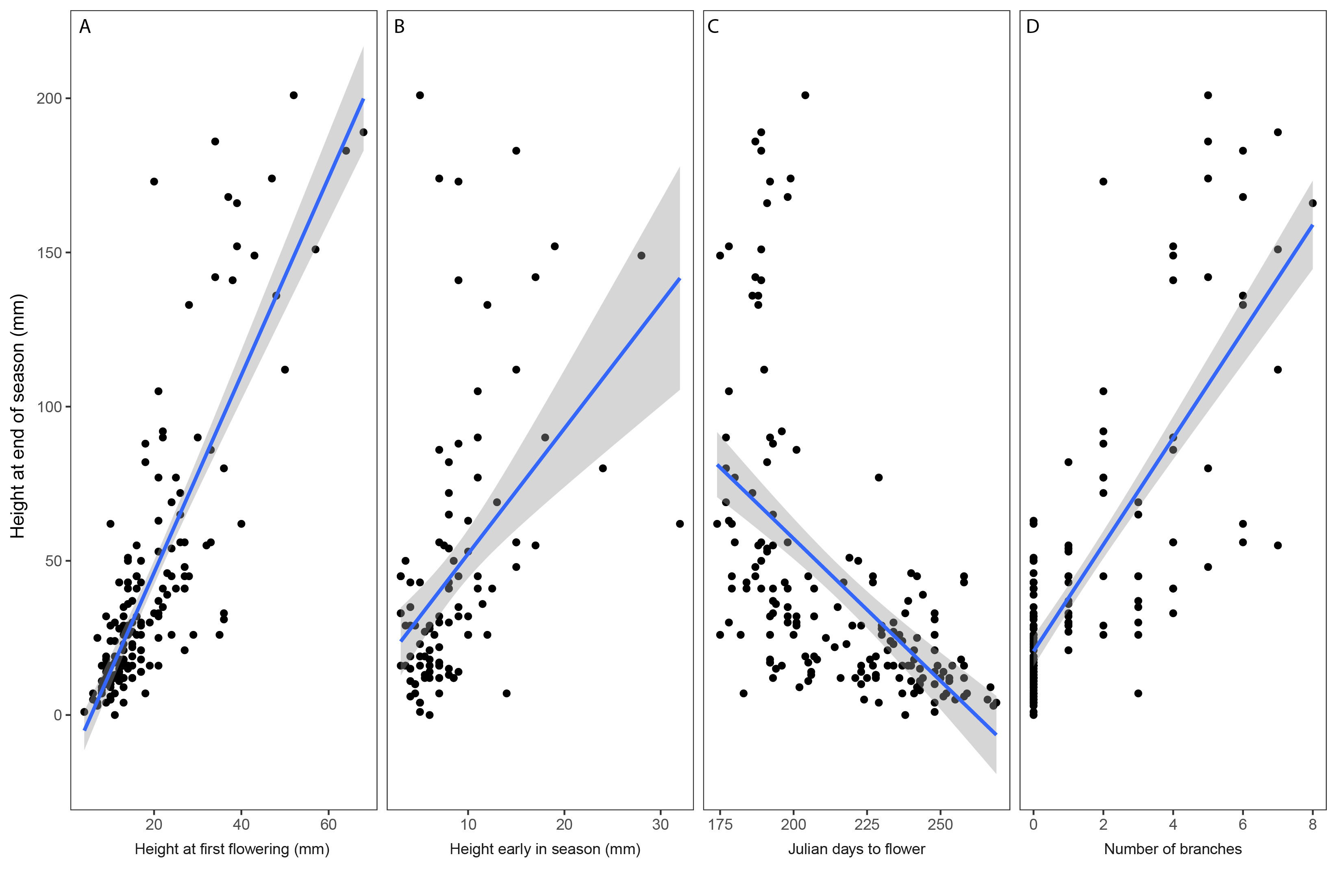

### Fig. S2

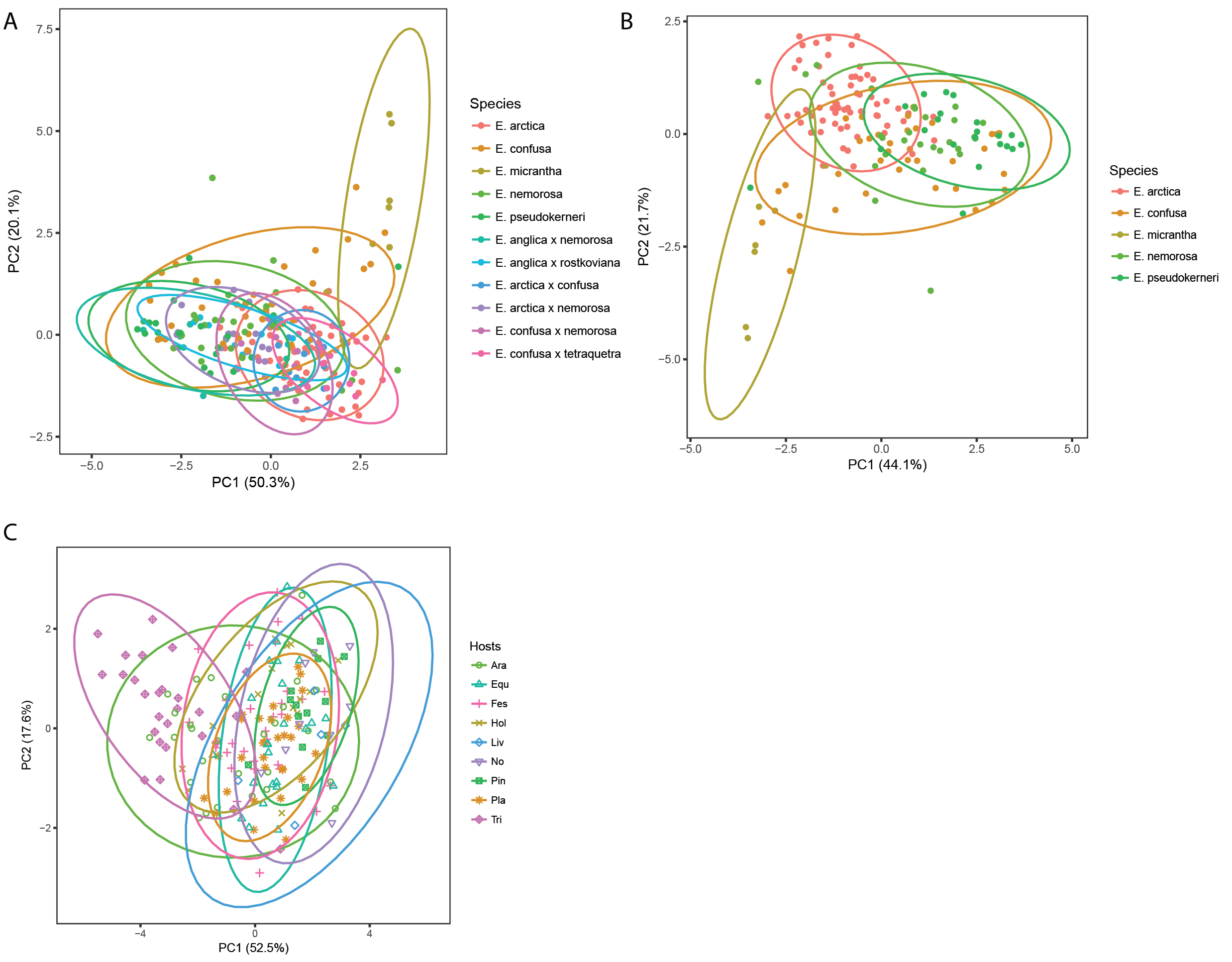
