## Supplementary material for "Life history evolution and phenotypic plasticity in parasitic eyebrights (*Euphrasia*, Orobanchaceae)": Table S1

**Table S1.** Host species used in the common garden experiment. Commercial seed stocks list the original collection where known.

| Common name | Species name | Family | Functional group (informal) | Seed source |
| --- | --- | --- | --- | --- |
| Thale cress | *Arabidopsis thaliana* | Brassicaceae | Herb | Laboratory stock |
| Field horsetail | *Equisetum arvense* | Equisetaceae | Fern | Wild collected in Edinburgh (GPS coordinates: 55.9679,-3.2129) |
| Red fescue | *Festuca rubra* | Poaceae | Grass | Commerical: Emorsgate seeds (Yorkshire + Dorset) |
| Yorkshire fog | *Holcus lanatus* | Poaceae | Grass | Commerical: Emorsgate seeds |
| Common liverwort | *Marchantia polymorpha* | Marchantiaceae | Bryophyte | Wild collected in Edinburgh (GPS coordinates: 55.9679,-3.2129) |
| Ribwort plantain | *Plantago lanceolata* | Plantaginaceae | Herb | Commerical: Emorsgate seeds (Somerset + Wiltshire) |
| Scots pine | *Pinus sylvestris* | Pinaceae | Tree | Commerical: Scotia Seeds |
| Red clover | *Trifolium pratense* | Fabaceae | Herb | Commerical: Emorsgate seeds (Yorkshire + Wiltshire) |
