## Supplementary material for "Life history evolution and phenotypic plasticity in parasitic eyebrights (*Euphrasia*, Orobanchaceae)": Table S2

**Table S2.** Collection details for *Euphrasia* species used in the common garden experiment. *Population also used in the multiple host phenotypic plasticity experiment.

| Collection number | Taxon | Locality | Latitude | Longitude | Collector |
| --- | --- | --- | --- | --- | --- |
| E4E0138 | *E. arctica* | Fintallick, Glen Ledock, Comrie, Perthshire | 56.41318 | -4.03085 | Dot Hall |
| E4E0144 | *E. arctica* | Balachuirn, Isle of Raasay | 57.38996 | -6.06877 | S.J. Bungard |
| E4E0032 | *E. arctica* | South Links, Burray, Orkney | 58.85275 | -2.88701 | John Crossley |
| E4E0139 | *E. arctica* | Dalreoch Farm, Enochdhu | 56.74199 | -3.53350 | Martin Robinson |
| E4E0049 | *E. arctica* | Ouaisne, Jersey | 49.17707 | -2.18293 | Anne Haden |
| E4E0247 | *E. arctica* | Elsdon. Newcastle upon Tyne | 55.22770 | -2.10234 | Stephanie Miles |
| NBer001* | *E. arctica* | North Berwick Glenn, East Lothian | 56.05696 | -2.70456 | Alex Twyford |
| E4E0038 | *E. confusa* | Oldbury, near Hartshill, Warwickshire | 52.55285 | -1.53980 | John & Monika Walton |
| E4E0114 | *E. confusa* | Trethew Mill, Bodmin, Cornwall | 50.39585 | -4.709558 | Rosemary Parslow |
| E4E0095 | *E. confusa* | North Anston Grassland, South Yorkshire | 53.34738 | -1.20803 | Graeme Coles |
| E4E0009 | *E. confusa* | Devil's Hole Blowout, Ravenmeols Local Nature Reserve, Merseyside | 53.54062 | -3.09041 | Philip H. Smith |
| E4E0188 | *E. micrantha* | Meall a Bathaich, Glen Garry, East Perthshire | 56.82082 | -4.182812 | Alistair Godfrey |
| E4E0064 | *E. nemorosa* | Castle Hill Local Nature Reserve, East Sussex | 50.7842 | 0.052719 | David Harris |
| E4E0069 | *E. nemorosa* | Meridian Business Park, Leicester | 52.60857 | -1.19809 | Geoffrey Hall |
| E4E0123 | *E. nemorosa* | Bloody Oaks Triangle, Tickercote, Rutland | 52.68950 | -0.56263 | Geoffrey Hall |
| E4E0029 | *E. pseudokerneri* | Levin Down, Sussex | 50.91346 | -0.74150 | Elizabeth Sturt |
| E4E0112 | *E. pseudokerneri* | Beeston Common, Norfolk | 52.93442 | 1.220071 | Francis Farrow |
| E4E0027 | *E. anglica* x *nemorosa* | West Dean Woods, Sussex | 50.93212 | -0.79735 | Elizabeth Sturt |
| E4E0016 | *E. anglica* x *rostkoviana* | Straduff Rathcabbin, Co. Tipperary | 53.11902 | -8.02454 | David Nash |
| E4E0033 | *E. arctica* x *confusa* | Nr Quoyorally, South Ronaldsay, Orkney | 58.75897 | -2.93473 | John Crossley |
| E4E0145 | *E. arctica* x *nemorosa* | Kylfakin, Wof, Skye | 57.26685 | -5.76042 | S.J. Bungard |
| E4E0021 | *E. arctica* x *nemorosa* | Dunamase, Co. Laois | 53.03153 | -7.21015 | David Nash |
| E4E0031 | *E. nemorosa* x *confusa* | Dolebury Fort, Somerset | 51.32605 | -2.79432 | C.W. Hurfurt |
| E4E0143 | *E. tetraquetra* x *confusa* | Ballyteige Burrow, Co Wexford, Ireland | 52.20268 | -6.64325 | Jim Hurley |
