## Supplementary material for "Life history evolution and phenotypic plasticity in parasitic eyebrights (*Euphrasia*, Orobanchaceae)": Table S3

**Table S3.** Summary of trait values for many *Euphrasia* species and hybrids grown on a clover host. Values are mean +/- one standard error. Length measurements are in mm.

| Taxon | Corolla length | Height | Internode ratio | Julian days to flower | Lower floral leaf teeth | Nodes to flower | Number of branches |
| --- | --- | --- | --- | --- | --- | --- | --- |
| *E. arctica* | 8.0 ± 0.2 | 82.9 ± 4.4 | 1.1 ± 0.1 | 195.2 ± 1.5 | 4.4 ± 0.1 | 8.6 ± 0.2 | 4.56 ± 0.2 |
| *E. confusa* | 6.9 ± 0.2 | 134.4 ± 7.2 | 1.6 ± 0.1 | 200.2 ± 2.4 | 5.3 ± 0.2 | 11.1 ± 0.4 | 7.26 ± 0.5 |
| *E. micrantha* | 5.6 ± 0.2 | 70.6 ± 8.1 | 3.0 ± 0.4 | * | 2.4 ± 0.3 | 8.3 ± 0.2 | 0.57 ± 0.4 |
| *E. nemorosa* | 7.7 ± 0.1 | 127.4 ± 8.1 | 1.4 ± 0.1 | 206.6 ± 1.7 | 5.1 ± 0.2 | 11.9 ± 0.5 | 7.67 ± 0.5 |
| *E. pseudokerneri* | 8.8 ± 0.4 | 176.4 ± 15.6 | 1.4 ± 0.1 | 205.1 ± 2.0 | 5.5 ± 0.2 | 13.2 ± 0.4 | 8.67 ± 0.6 |
| *E. anglica* x *nemorosa* | 9.1 ± 0.5 | 148.1 ± 11.8 | 1.4 ± 0.1 | 195.7 ± 1.9 | 6.0 ± 0.3 | 12.0 ± 0.6 | 10.00 ± 1.0 |
| *E. anglica* x *rostkoviana* | 7.9 ± 0.2 | 122.6 ± 8.3 | 1.3 ± 0.1 | 192.3 ± 12.3 | 5.9 ± 0.3 | 10.6 ± 0.5 | 7.44 ± 0.7 |
| *E. arctica* x *confusa* | 9.5 ± 0.2 | 100.3 ± 4.3 | 1.4 ± 0.1 | 193.4 ± 3.2 | 3.8 ± 0.1 | 7.8 ± 0.3 | 5.70 ± 0.4 |
| *E. arctica* x *nemorosa* | 8.0 ± 0.2 | 132.2 ± 14.5 | 1.3 ± 0.1 | 205.3 ± 2.4 | 6.0 ± 0.3 | 11.3 ± 0.4 | 6.50 ± 0.4 |
| *E. confusa* x *nemorosa* | 7.9 ± 0.2 | 92.5 ± 5.9 | 1.0 ± 0.1 | 199.3 ± 2.8 | 5.1 ± 0.2 | 9.8 ± 0.3 | 7.00 ± 0.5 |
| *E. confusa* x *tetraquetra* | 7.2 ± 0.2 | 57.4 ± 5.8 | 0.7 ± 0.1 | 194.1 ± 2.7 | 4.2 ± 0.2 | 7.6 ± 0.4 | 4.00 ± 0.3 |

*Date of first flower not recorded.
