## Supplementary material for "Life history evolution and phenotypic plasticity in parasitic eyebrights (*Euphrasia*, Orobanchaceae)": Table S4

| (a) Species differences (including hybrids) |  |  |  |  |  |
| --- | --- | --- | --- | --- | --- |
|  | PC1 | PC2 | PC3 | PC4 | PC5 |
| Branches | 0.180 | 0.066 | 0.028 | 0.116 | 0.174 |
| Corolla length | 0.137 | 0.026 | 0.267 | 0.335 | 0.006 |
| Height | 0.175 | 0.132 | 0.007 | 0.104 | 0.161 |
| Internode ratio | 0.085 | 0.262 | 0.331 | 0.146 | 0.116 |
| Julian days to flower | 0.156 | 0.201 | 0.042 | 0.047 | 0.208 |
| Leaf teeth | 0.176 | 0.012 | 0.089 | 0.130 | 0.182 |
| Nodes to flower | 0.091 | 0.302 | 0.236 | 0.122 | 0.153 |
| Standard deviation | 1.917 | 1.109 | 0.897 | 0.731 | 0.621 |
| Proportion of variance | 0.525 | 0.176 | 0.115 | 0.076 | 0.055 |
| (b) Species differences (excluding hybrids) |  |  |  |  |  |
|  | PC1 | PC2 | PC3 | PC4 | PC5 |
| Branches | 0.226 | 0.024 | 0.096 | 0.233 | 0.017 |
| Corolla length | 0.100 | 0.269 | 0.361 | 0.082 | 0.141 |
| Height | 0.214 | 0.128 | 0.151 | 0.063 | 0.367 |
| Internode ratio | 0.029 | 0.434 | 0.202 | 0.000 | 0.171 |
| Leaf teeth | 0.214 | 0.032 | 0.064 | 0.424 | 0.159 |
| Nodes to flower | 0.217 | 0.113 | 0.125 | 0.198 | 0.145 |
| Standard deviation | 1.780 | 1.111 | 0.932 | 0.612 | 0.433 |
| Proportion of variance | 0.528 | 0.206 | 0.145 | 0.062 | 0.031 |
| (c) Phenotypic plasticity |  |  |  |  |  |
|  | PC1 | PC2 | PC3 | PC4 | PC5 |
| Branches | 0.180 | 0.066 | 0.028 | 0.116 | 0.174 |
| Corolla length | 0.137 | 0.026 | 0.267 | 0.335 | 0.006 |
| Height | 0.175 | 0.132 | 0.007 | 0.104 | 0.161 |
| Internode ratio | 0.085 | 0.262 | 0.331 | 0.146 | 0.116 |
| Julian days to flower | 0.156 | 0.201 | 0.042 | 0.047 | 0.208 |
| Leaf teeth | 0.176 | 0.012 | 0.089 | 0.130 | 0.182 |
| Nodes to flower | 0.091 | 0.302 | 0.236 | 0.122 | 0.153 |
| Standard deviation | 1.917 | 1.109 | 0.897 | 0.731 | 0.621 |
| Proportion of variance | 0.525 | 0.176 | 0.115 | 0.076 | 0.055 |
