## Supplementary material for "Life history evolution and phenotypic plasticity in parasitic eyebrights (*Euphrasia*, Orobanchaceae)": Table S5

**Table S5.** Summary of trait values for *Euphrasia arctica* grown on many different hosts. Values are mean +/- one standard error. Length and height measurements are in mm.

| Host | Early season | | At first flowering | | | | | | |  | End of season | |
| --- | --- | --- | --- | --- | --- | --- | --- | --- | --- | --- | --- | --- |
|  | Height | Corolla length | | Height | Internode ratio | Julian days to flower | Lower floral leaf teeth | Nodes to flower | Number of branches | | | Height |
| *A. thaliana* | 12.78 ± 1.47 | 6.09 ± 0.28 | | 19.17 ± 1.60 | 2.38 ± 0.13 | 201.57 ± 4.3 | 3.17 ± 0.14 | 8.83 ± 0.28 | 2.10 ± 0.39 | | | 40.54 ± 4.17 |
| *E. arvense* | 6.07 ± 0.55 | 5.94 ± 0.31 | | 15.12 ± 1.08 | 2.62 ± 0.18 | 215.25 ± 4.55 | 2.36 ± 0.11 | 9.28 ± 0.30 | 0.36 ± 0.13 | | | 31.91 ± 4.24 |
| *F. rubra* | 6.69 ± 0.61 | 6.29 ± 0.13 | | 19.45 ± 1.37 | 2.59 ± 0.15 | 216.52 ± 4.39 | 2.84 ± 0.16 | 9.58 ± 0.25 | 0.84 ± 0.25 | | | 46.33 ± 3.18 |
| *H. lanatus* | 7.10 ± 1.64 | 6.30 ± 0.14 | | 16.00 ± 1.59 | 2.36 ± 0.19 | 224.47 ± 6.95 | 2.47 ± 0.19 | 9.82 ± 0.35 | 0.82 ± 0.35 | | | 46.13 ± 6.02 |
| *M. polymorpha* | 6.33 ± 0.84 | 5.50 ± 0.42 | | 9.56 ± 1.30 | 2.90 ± 0.40 | 222.60 ± 16.9 | 1.67 ± 0.29 | 9.67 ± 0.47 | 0 | | | 21.80 ± 7.00 |
| No host | 5.92 ± 0.42 | 5.25 ± 0.24 | | 11.18 ± 1.09 | 2.75 ± 0.23 | 241.33 ± 7.85 | 1.91 ± 0.25 | 9.91 ± 0.48 | 0 | | | 30.13 ± 7.83 |
| *P. lanceolata* | 7.47 ± 0.61 | 6.13 ± 0.10 | | 14.13 ± 0.75 | 2.84 ± 0.12 | 211.19 ± 3.73 | 2.90 ± 0.11 | 10.43 ± 0.27 | 0.39 ± 0.14 | | | 35.17 ± 3.65 |
| *P. sylvestris* | 6.25 ± 0.71 | 5.73 ± 0.25 | | 12.21 ± 1.31 | 2.90 ± 0.22 | 233.77 ± 6.14 | 1.93 ± 0.16 | 9.23 ± 0.27 | 0 | | | 29.17 ± 7.07 |
| *T. pratense* | 12.91 ± 1.71 | 7.35 ± 0.17 | | 39.44 ± 2.64 | 2.10 ± 0.17 | 189.81 ± 2.00 | 3.89 ± 0.14 | 8.67 ± 0.21 | 4.74 ± 0.38 | | | 20.08 ± 3.59 |
